# Multi-omics insights into the response of *Aspergillus parasiticus* to long-chain alkanes in relation to polyethylene degradation

**DOI:** 10.1101/2024.12.16.628742

**Authors:** Romanos Siaperas, George Taxeidis, Efstratios Nikolaivits, Evangelos Topakas

## Abstract

Plastic pollution presents a global challenge, with polyethylene (PE) being among the most persistent plastics due to its durability and environmental resilience. In this study, we employed a multi-omics approach to study the ability of *Aspergillus parasiticus* MM36, an isolate derived from *Tenebrio molitor* intestines, to metabolize long-chain alkanes (lcAlk) and secrete enzymes able to modify PE. The fungus was grown with hexadecane (C16) or a mixture of lcAlk (C24 to C36) as carbon sources and culture supernatants were tested daily for their ability to modify PE. Proteomic analysis identified induced oxidases potentially involved in lcAlk and PE functionalization. Key enzymes include multicopper oxidases, peroxidases, an unspecific peroxygenase and FAD-dependent monooxygenases. Surfactant proteins facilitating enzymatic and cellular interaction with hydrophobic PE and lcAlk, such as one hydrophobin, three hydrophobic surface-binding proteins (HsbA) and one cerato platanin, were present in all secretomes. Transcriptomic analysis comparing lcAlk to C16 cultures highlighted the enrichment of oxidoreductase activities and carboxylic acid metabolism in both lcAlk incubation days, with transmembrane transporters and transferases predominating on day 2 and biosynthetic processes on day 3. In C16 cultures, hydrolytic enzymes, including esterases, were upregulated alongside Baeyer-Villiger monooxygenases, suggesting a shift toward sub-terminal hydroxylation. Integrating transcriptomic and secretomic data, we propose a mechanism for lcAlk assimilation by *A. parasiticus* MM36, involving extracellular oxyfunctionalization, hydrocarbon uptake via surface-modifying proteins and channeling through membrane transporters for energy consumption and biosynthetic processes. This study provides insights into fungal mechanisms for alkane metabolism and highlights their relevance to plastic degradation.

**Importance:** Plastic pollution presents a global challenge to marine life and human health, with polyethylene (PE) being among the most persistent plastics due to its durability and environmental resilience. Hydroxylation is regarded as the initial step of PE degradation, similar to alkane oxidation, making alkane-degrading microbes a promising source of plastic degraders. In this study, we used a multi-omics approach to investigate the ability of *Aspergillus parasiticus* MM36 to metabolize long-chain alkanes and secrete enzymes that modify PE. Proteomic analysis of the secretomes identified key oxidases and biosurfactants that enable the fungus to interact with and transform hydrophobic substrates like PE. Transcriptomic analysis further revealed biological processes involved in alkane assimilation and metabolism. By integrating these insights, we propose a mechanism for fungal alkane metabolism and highlight its relevance to plastic biodegradation. This work advances our understanding of fungal contributions to addressing hydrocarbon and plastic pollution.

## 1. Introduction

Plastic pollution poses a global challenge with synthetic polymers accumulating in the environment since their mass production began in the 1950s. Plastics are produced by polymerization of monomers derived from oil or gas and enhanced with chemical additives (Thompson et al., 2009). Among these, polyethylene (PE) stands out as one of the most prevalent plastics due to its versatility, durability and cost effectiveness accounting for over 22% of European plastic production in 2022 (Plastics Europe, 2024). PE is a thermoplastic polyolefin, with ethylene being the single monomer unit. Its high molecular weight, hydrophobic nature and lack of functional groups render PE exceptionally resistant, with discarded PE items persisting for decades to centuries (Chamas et al., 2020). Improper disposal and the subsequent formation of microplastics pose risks to marine life and human health through bioaccumulation (Filella et al., 2021).

Microbial degradation of plastics has emerged as a promising avenue for tackling plastic pollution. Many microorganisms, including bacteria and fungi, can assimilate carbon derived from synthetic petrochemical polymers by repurposing enzyme machinery originally evolved for degrading structurally similar natural polymers (Taxeidis et al., 2024; Zerva et al., 2021). Alkane-degrading microbes have shown potential for PE degradation. Bacteria such as *Rhodococcus* and *Alcanivorax* can grow in the presence of PE (Gravouil et al., 2017; Zampolli et al., 2021). Similarly, fungal alkane-degrading genera such as *Penicillium* and *Aspergillus* (Al-Hawash et al., 2018; Velez et al., 2020) have been reported to act on PE (Yamada-Onodera et al., 2001; Zhang, Gao, et al., 2020). Furthermore, the ability of insect larvae, particularly *Tenebrio molitor* and its gut microbiota, to degrade PE has been widely studied (Pivato et al., 2022). These larvae naturally consume beeswax, a mixture rich in alkanes, alkenes, fatty acids, and esters (Maia & Nunes, 2013). A recent study highlighted the dominance of the *Aspergillaceae* family in the gut of *T. molitor* after feeding with polypropylene, a polymer classified under polyolefins alongside PE (Y. Wang et al., 2023).

Although PE biodegradation has been demonstrated, its underlying mechanisms remain poorly understood (Restrepo-Flórez et al., 2014). The only reported microbial PE-degrading enzymes are three laccases from *Rhodococcus (Tao et al., 2023; Zampolli et al., 2023)*. Like alkane degradation, PE biodegradation is hypothesized to begin with hydroxylation either in-chain or at the termini (Jin et al., 2023). Alkane metabolism has been extensively explored at the omics level in bacteria and yeasts, but studies on filamentous fungi remain sparse. Only one transcriptomic study has examined the growth of a *Penicillium* sp. on hexadecane (C16) and hexadecene (Velez et al., 2020). For PE biodegradation, omics analyses have predominantly focused on hydrocarbonoclastic bacteria like *Rhodococcus* and *Alcanivorax (Gravouil et al., 2017; Tao et al., 2023; Zadjelovic et al., 2022; Zampolli et al., 2021)*, employing transcriptomics or proteomics. Among fungi, there are only two relevant studies: a transcriptomic analysis of the marine fungus *Alternaria alternata* FB1 in the presence of PE (Gao et al., 2022), and a proteomics study on the yeast *Yarrowia lipolytica* for upcycling of depolymerized PE (Caleb et al., 2023).

Alkane chain length influences their hydrophobicity and bioavailability (Rojo et al., 2008). Gram-negative bacteria encode transporters able to transfer alkanes up to C38 through their outer membrane to process them intracellularly (Liu et al., 2022). Fungi accumulate short and mid-chain alkanes like C16 (Al-Hawash et al., 2018; Li et al., 2021), but longer-chain alkanes likely require functionalization or chain scission at the extracellular space —a necessary step for PE as well (Zadjelovic et al., 2022).

Previous research demonstrated that *Aspergillus parasiticus* MM36, isolated from *T. molitor* intestines, can grow using a mixture of long-chain alkanes (lcAlk), ranging from C24 to C36, as the sole carbon source. Additionally, its lcAlk-induced secretome can cause structural modifications on PE (Taxeidis et al., 2023). To gain deeper insights into this strain’s capabilities, we employed a multi-omics approach, starting with genome sequencing, assembly and annotation. We performed RNAseq to investigate the molecular response during lcAlk assimilation. Secretome analysis with LC-MS/MS provided further insights into the enzymes involved in lcAlk and PE modification. This offers a comprehensive view of alkane metabolism by filamentous fungi, highlighting its relevance to plastic degradation.

## 2. Results and Discussion

### 2.1 Genomic analysis of newly isolated Aspergillus parasiticus MM36

A. *parasiticus* MM36, isolated from mealworm (*Tenebrio molitor*) intestines, can grow using lcAlk as the sole carbon source and its lcAlk-induced secretome causes structural modifications in PE (Taxeidis et al., 2023). To study the genetic features underlying this capability and to facilitate subsequent transcriptomic and proteomic analyses, we first performed whole genome Illumina sequencing and annotation. The assembled genome comprises 321 scaffolds totaling 39.8 Mb. Its predicted proteome includes 13844 proteins with a 99.6 % BUSCO completeness, comparable to the reference proteome of *A. parasiticus* CBS 117618 (13752 proteins, 98.3 % completeness) (Kjærbølling et al., 2020). The repeat content in the *A. parasiticus* MM36 genome, 2.90 % as determined by RepeatMasker, is consistent with the 2-3 % average of the *Aspergillaceae* family (de Vries et al., 2017). There are 1409 transport-related genes, identified through the tcdb database (Saier Jr et al., 2021), with electrochemical potential-driven transporters, particularly the major facilitator superfamily (MFS), being the most abundant family. This family shows the highest level of expansion in the *Eurotiomyces* class (de Vries et al., 2017).

Α multilocus phylogenetic analysis was conducted to confirm the species classification, as previous ITS sequence-based attempts were inconclusive. Two hundred (200) single-copy orthologs were included from 29 genomes within the *A. Flavi* section and two outgroup species. The methodology for ortholog generation and alignment was based on the work of Steenwyk et al (Steenwyk et al., 2024) and the genomes were primarily sourced from the study of Kjærbølling et al. (Kjærbølling et al., 2020). Our results confirm MM36 is part of the A. *parasiticus* species forming a well-supported clade of 100 ultrafast bootstrap and SH-like approximate likelihood ratio test with *A. parasiticus* CBS 117618 (Fig. S1). The moderate gene concordance factor (42.1 %) of this clade indicates limited signal for many single-gene trees, however 81.2 % of the informative sites also supported this clade (Minh, Hahn, et al., 2020).

### 2.2 Determining the optimal time points for multi-omics analyses

To select optimal time points for transcriptomics and proteomics analyses, we assessed the ability of *A. parasiticus* MM36 secretomes to modify LDPE structure over time. Secretomes from cultures grown on lcAlk for 2 days (lcAlk_2d) caused significant changes in the LDPE molecular fingerprint. Similar modifications were observed after growth in C16 and lcAlk for 3 days (C16_3d and lcAlk_3d).

To better interpret and compare these results, intensity ratios between the characteristic peaks of LDPE and newly formed peaks were calculated from the ATR-FTIR spectra. Specifically, the peaks at 2919 cm⁻¹ and 1462 cm⁻¹, corresponding to CH₃ stretching vibrations and C-H bending deformations respectively, were selected due to their typical presence in untreated LDPE (Kovács et al., 2021). Concurrently, the peak at 1210 cm⁻¹ that corresponds to C–O stretching, an indicative sign of acyl groups formation, was selected as it can be correlated with LDPE functionalization (Fig. 1).

**Figure 1:**
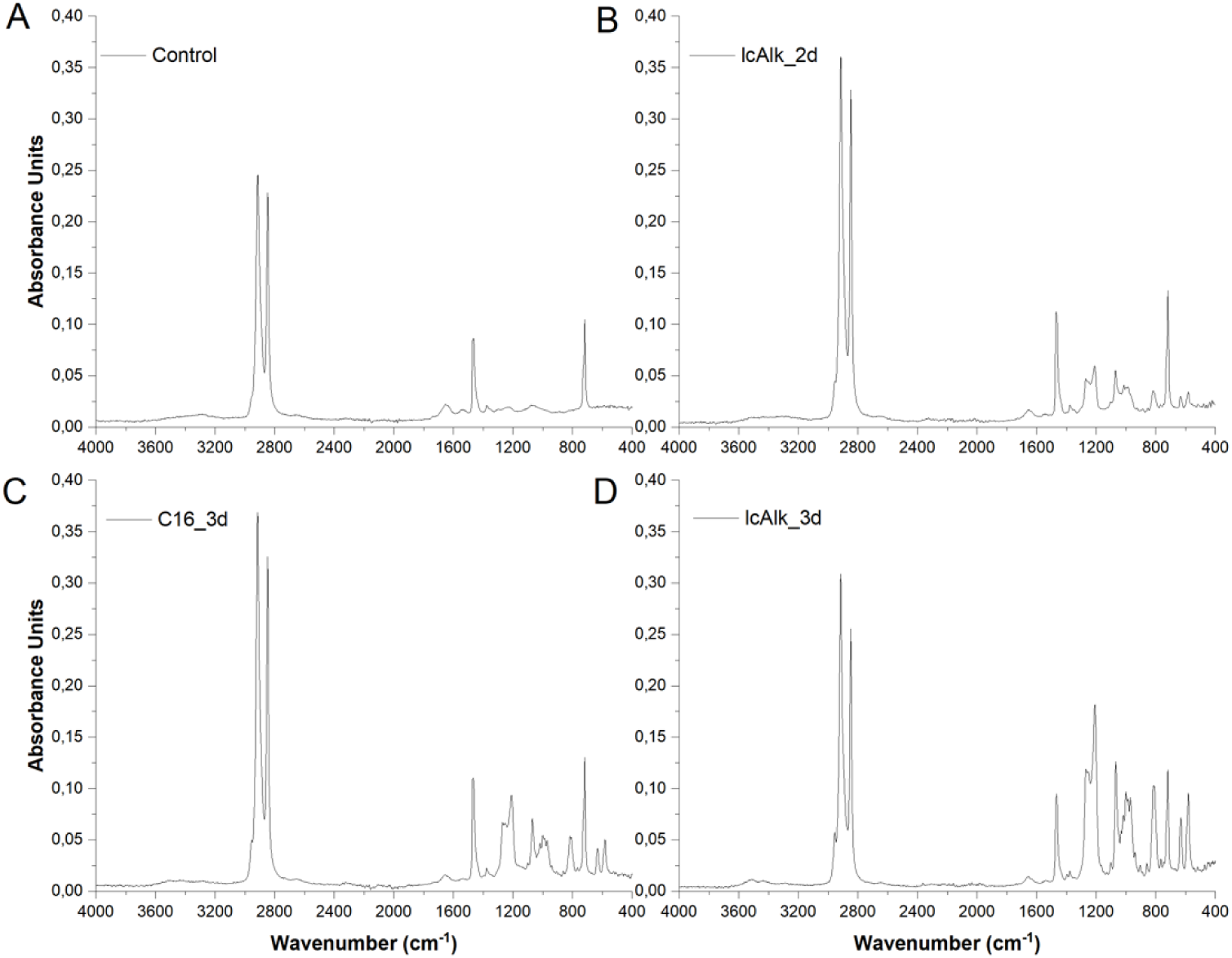
ATR-FTIR spectrum of virgin LDPE powder (A) and LDPE powder after treatment with the secretome of *A. parasiticus* MM36, when grown in presence of lcAlk for 2 days (B), and C16 (C) or lcAlk (D) for 3 days, respectively.

As presented in Table 1, the calculated ratios between the characteristic peaks at 1462 cm⁻¹ and 2919 cm⁻¹ remain nearly constant under all tested conditions fluctuating from 0.30-0.37. At the same time, the ratio between the peaks at 1210 cm⁻¹ and 2919 cm⁻¹ increases progressively, reaching the maximum value after treating LDPE with the lcAlk_3d secretome (Table 1). Moreover, this specific intensity ratio is 2.3-fold higher when compared to the C16_3d control condition, marking the highest observed value between lcAlk and C16 during this time-course experiment. This significant difference suggests that the lcAlk_3d secretome can be possibly enriched with different and/or more abundant LDPE-converting enzymes. Another noteworthy point is that the lcAlk_2d secretome, which was also selected for further analysis, exhibited significantly lower intensity ratios between the peaks at 1210 cm⁻¹ and 2919 cm⁻¹ compared to the lcAlk_3d secretome (3.5-fold lower intensity ratio), suggesting that the microorganism has been better adapted towards lcAlk breakdown after 3 days. These preliminary findings indicate that lcAlk_3d secretome likely contains proteins directly associated with efficient LDPE functionalization. This contrasts with the lcAlk_2d and C16_3d secretomes, which were utilized as comparative proxies in the subsequent differential analysis.

**Table 1:** Calculated intensity ratios between characteristic PE peaks and newly formed on ATR-FTIR spectra.

| Sample | Intensity ratio | Intensity ratio |
| --- | --- | --- |
|  | 1462 cm<sup>-1</sup> / 2919 cm<sup>-1</sup> | 1210 cm<sup>-1</sup> / 2919 cm<sup>-1</sup> |
| Control | 0.35 | 0.07 |
| lcAlk_1d | 0.36 | 0.11 |
| **lcAlk_2d** | 0.31 | 0.17 |
| **lcAlk_3d** | 0.31 | 0.59 |
| lcAlk_4d | 0.31 | 0.39 |
| C16_1d | 0.37 | 0.13 |
| C16_2d | 0.35 | 0.08 |
| **C16_3d** | 0.30 | 0.26 |
| C16_4d | 0.31 | 0.48 |

### 3.3 Secretomic analysis reveals strong substrate and time-dependent variability

We analysed the secretomes of *A. parasiticus* MM36 grown under three selected conditions (lcAlk_2d, lcAlk_3d and C16_3d) using label-free data-dependent LC-MS/MS proteomics in triplicates to identify enzymes potentially acting on PE and facilitating lcAlk assimilation. The assimilation of lower molecular weight alkanes from the water phase is feasible due to their solubility (Rojo, 2009). In contrast, lcAlk assimilation requires extracellular oxyfunctionalization (Zadjelovic et al., 2022).

A total of 20.7 % of the total MS/MS spectra were successfully assigned to peptides in the protein database, identifying 5,230 peptides that aggregated into 1,096 proteins. On average, 25 % of the spectra analysed in an MS experiment can be identified, however this percentage is expected to drop when analysing the secretome instead of the whole proteome (Griss et al., 2016) as observed in Brana et. al. (Pantelic et al., 2024). After applying a filter for proteins present in at least two replicates of any condition, 722 proteins were reliably quantified, with 34.6 % predicted to be extracellular compared to 10.0 % of the total proteome. The predominant enzyme group was oxidoreductases (140 proteins), followed by 74 proteins of unknown function. Forty-three of these uncharacterized proteins along with 38 other proteins in the secretomes are shorter than 300 amino acids and are classified as small secreted proteins (SSPs). The primary biological role of many SPPs in host-associated fungi like *A. parasiticus* MM36 is the interaction with the living host (Feldman et al., 2020).

Secretome profiles showed strong substrate- and time-dependent variability, with the lkAlk_3d condition displaying the richest secretome encompassing almost all quantified proteins. In contrast, the lcAlk_2d condition exhibited the fewest proteins, with only half of the total proteins observed across all conditions. Most differentially abundant proteins were induced in the lcAlk_3d secretomes, with 154 differing from lcAlk_2d and 7 from C16_3d, 6 of which were common. This result aligns with the ATR-FTIR analysis, which showed maximal PE modification by lcAlk_3d secretomes and activity in lcAlk_2d secretomes.

Differential analysis highlighted a predominance of hydrolytic enzymes, including 15 peptidases, 16 glycoside hydrolases, 7 esterases, three phosphatases, one amidase, and one cysteinase. Six glycoside hydrolases contain carbohydrate-binding modules, and additional induced proteins included three carbohydrate-binding lectins and one LysM (lysin motif) domain-containing protein.

### 2.4 Identification of potential plastizymes in Aspergillus parasiticus MM36 secretomes

We sought to identify enzymes potentially capable of oxidizing PE within the secretomes of *A. parasiticus* MM36. Multicopper oxidases (MCOs) constitute the AA1 CAZy family and contain laccases, ferroxidases and *Ascomycota* laccase-like enzymes (Levasseur et al., 2013). To date, two MCOs from *Rhodococcus opacus* R7 are the only isolated enzymes shown to oxidize PE and release a variety of alkanes and oxygenated hydrocarbons (Zampolli et al., 2023). Additionally, an MCO from *Parvibaculum lavamentivorans* degrades C19-C25 alkanes with a yield of more than 90 % (Diefenbach et al., 2024). Four MCOs were quantified in the secretomes of *A. parasiticus* MM36. The AA1_3 laccase-like RU639_012907 was induced in lcAlk_3d compared to both lcAlk_2d and C16_3d. Another MCO (RU639_007412) was induced compared to lcAlk_2d and shares 94.6 % amino acid identity with AFLA_006190, one of the two MCOs upregulated during the degradation of PE by *A. flavus* PEDX3 (Fig. 2) (Zhang, Gao, et al., 2020).

**Figure 2:**
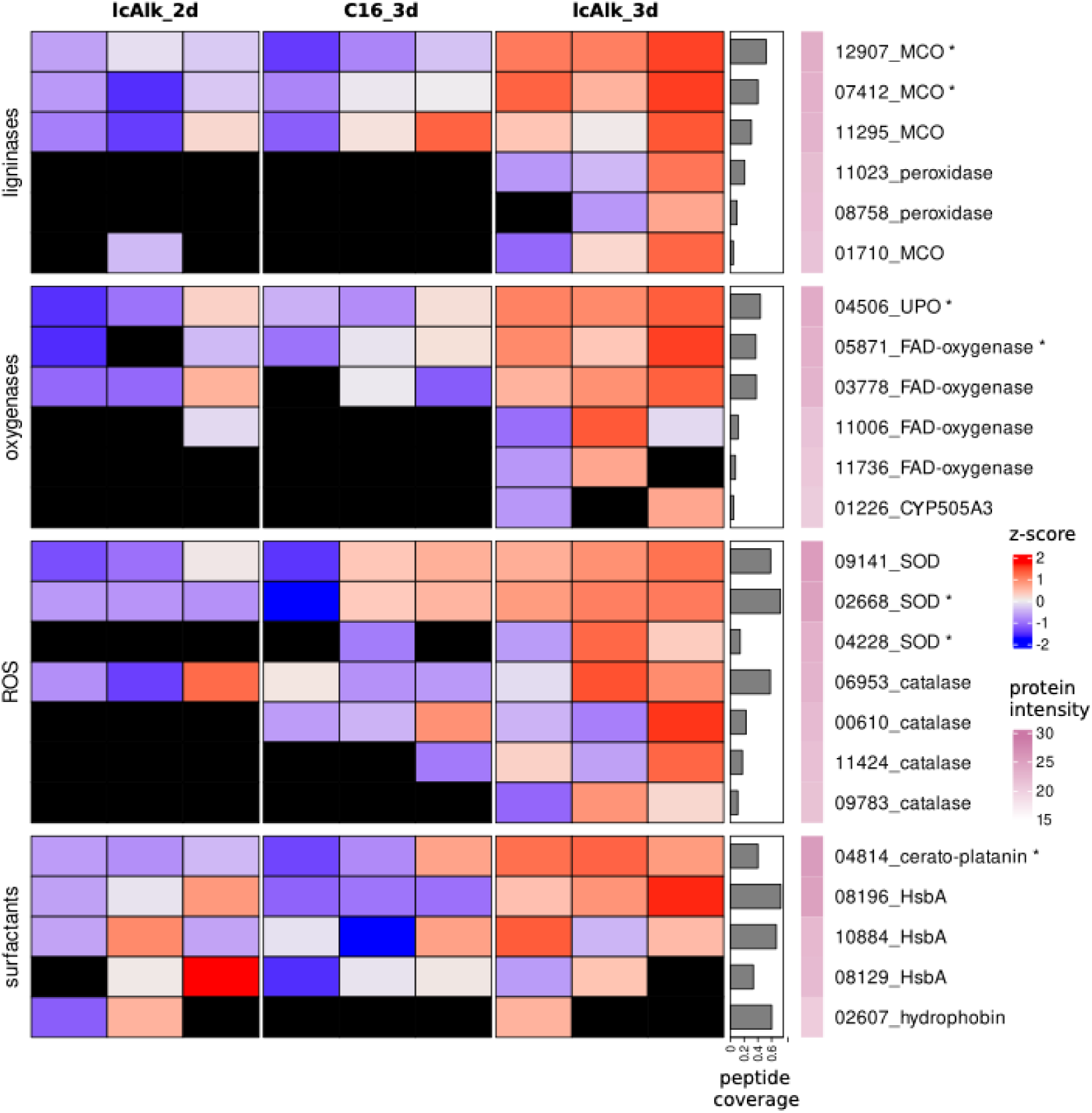
Heatmap of z-scores of protein intensities for selected proteins; black color means not detected. Proteins differentially abundant in lcAlk_3d compared to lcAlk_2d are denoted with an asterisk. peptide coverage: Number of detected peptides divided by number of theoretical tryptic peptides. protein intensity: log2 of average protein intensity in lcAlk_3d. ROS: reactive oxygen species; MCO: multicopper oxidase; UPO: unspecific peroxygenase; SOD: superoxide dismutase; HsbA: hydrophobic surface-binding protein A.

Unspecific peroxygenases (UPOs) are extracellular heme proteins that catalyse various C-H oxyfunctionalization reactions using hydrogen peroxide as an oxygen donor (Faiza et al., 2019). UPO1 from *Agrocybe aegerita* hydroxylates linear alkanes up to C16 but is inactive on longer chains (Peter et al., 2011). MroUPO from the white-rot fungus *Marasmius rotula* initiates a cascade of mono- and di-terminal oxygenation reactions of dodecane (C12) and tetradecane (C14) to the corresponding carboxylic acids (Olmedo et al., 2016). UPOs are classified in InterPro as members of the chloroperoxidase protein family (IPR000028) (Paysan-Lafosse et al., 2023). The genome of *A. parasiticus* MM36 encodes five extracellular chloroperoxidases; among these only RU639_007575 was detected in our experiment and was induced in lcAlk_3d secretomes. RU639_007575 is a short UPO of 27.5 kDa that contains the characteristic PCP (Proline-Cysteine-Proline) amino acid motif for heme-binding, as well as the EHD (Glutamate-Histidine-Aspartate) motif. The latter is a conserved signature for UPOs like MroUPO (Faiza et al., 2019). The most studied lcAlk oxidases, able to hydroxylate alkanes up to C36 to their primary alcohols, are the bacterial flavin-dependent monooxygenases named LadA and almA (Feng et al., 2007; W. Wang & Shao, 2014). LadA belongs to the luciferase family (IPR016215), while AlmA is part of the FAD-binding monooxygenase family (IPR051820). However, enzymes from these families in *A. parasiticus* MM36 are intracellular and were not detected in the secretomes. In contrast, four flavin-dependent monooxygenases belonging to a distinct FAD-binding superfamily (IPR036318) were identified in the secretomes, with RU639_005614 being differentially abundant in lcAlk_3d compared to lcAlk_2d (Fig. 2). Another monooxygenase of interest is CYP505A3, secreted exclusively in lcAlk_3d. Members of the CYP505 family of the cytochrome P450 monooxygenases are considered sub-terminal fatty acid hydroxylases, while CYP505E3 from *A. terreus* catalyses also the regioselective in-chain hydroxylation of C10-C16 n-alkanes (Maseme et al., 2020).

Alkane hydroxylases overcome the low chemical reactivity of the alkanes by generating reactive oxygen species (ROS) (Rojo, 2009), a strategy also proposed for PE oxidation (Zadjelovic et al., 2022). ROS could also assist the action of the mentioned secreted monooxygenases. Additionally, ROS activate oxidative enzymes such as peroxidases. A bacterial peroxidase degrades polystyrene, an aromatic polymer with a C-C backbone similar to PE that generates water soluble products in the presence of H_2_O_2_ (Nakamiya et al., 1997). The lcAlk_3d secretomes contain two lignin peroxidases undetected in other conditions, a member of the AA2 CAZy family, and a DyP-type peroxidase. Interestingly, three superoxide dismutases were also induced in lcAlk_3d compared to lcAlk_2d (Fig. 2). These enzymes help control ROS levels, protecting the fungus from oxidative stress and preventing enzyme inhibition (Zhang, Hao, et al., 2020). While catalases, which also control ROS levels, were quantified at similar levels across all conditions, the induction of a superoxide dismutase rather than catalases has also been observed in *Alcanivorax* sp. 24 when exposed to pristine LDPE (Zadjelovic et al., 2022). Additionally, one thioredoxin, a protein associated with oxidative stress, and one thioredoxin reductase were detected only in the secretomes of lcAlk_3d.

Beyond oxidative enzymes, proteins involved in hydrocarbon uptake in fungi include those that modify the hydrophobicity of the cell surface or the hydrocarbon itself to enable cell adhesion. Hydrophobins promote sorption by increasing the cell surface hydrophobicity and constitute the IPR001338 Interpro family. This is exemplified in the fungus *Paecilomyces lilacinus*, which grows on C16 producing hydrophobins in solid-state cultures (Vigueras et al., 2014). The genome of *A. parasiticus* MM36 encodes three extracellular hydrophobins, but only the hydrophobin RU639_002607, an ortholog of RolA from *Aspergillus oryzae,* was detected by MS in two replicates of lcAlk_2d and one replicate of lcAlk_3d (Fig. 2). RolA is a hydrophobin that is known in literature to enhance PET degradation underpinning its potential for PE modification (Puspitasari et al., 2021).

Additionally, the secretomes contained three members of Hydrophobic surface-binding protein A (HsbA) family (IPR021054), with similar abundance across conditions (Fig. 2). One of these contains a GPI-anchor that keeps it attached to the cell wall. A HsbA from the pathogen *Penicillium marneffei* binds long-chain fatty acids and stores them in the cell wall for nutrient starvation resilience (Liao et al., 2010). RU639_010884 is orthologous to a HsbA from *A. oryzae* that binds hydrophobic polybutylene succinate-co-adipate (PBSA) promoting its degradation via a cutinase (Shinsaku et al., 2006). Cerato platanins are another protein family with surfactant properties that reduce the surface hydrophobicity of plastics (Renwei et al., 2020). *A. parasiticus* genome encodes two cerato platanins, one of which was constitutively secreted across all samples.

### 2.5 Transcriptomic profiling during alkane assimilation

To study the response of *A. parasiticus* MM36 to lcAlk and C16, and to identify the biological processes involved in their assimilation, we conducted RNA-seq at the same three conditions as secretomics analysis. After filtering out low-count genes, 11,964 genes were retained for differential expression analysis and the subsequent enrichment analyses. Overall, gene expression changes between the lcAlk timepoints were modest, with fewer than 250 genes differentially expressed, contrasting with the more pronounced variations observed at the secretome level. Conversely, numerous genes displayed differential expression when comparing both lcAlk timepoints to C16_3d, with approximately 1,000 genes upregulated in lcAlk conditions and nearly double in C16_3d.

Genes were mapped to gene ontology (GO) terms using InterProScan, and differentially expressed genes were tested for the enrichment of biological processes and molecular functions with GOATOOLS (Klopfenstein et al., 2018). Analysis comparing lcAlk_2d with C16_3d revealed an enrichment of genes associated with transmembrane transport processes, predominantly involving the Major Facilitator Superfamily (MFS) and ATP-binding Cassette (ABC) transporter families (Fig. 3). Similarly, the majority of *Rhodococcus ruber* transporters upregulated in the presence of PE were MFS and ABC transporters (Gravouil et al., 2017). Similarly, in *Yarrowia lipolytica*, an ABC transporter is involved in the uptake of C16 (Thevenieau et al., 2007).

**Figure 3:**
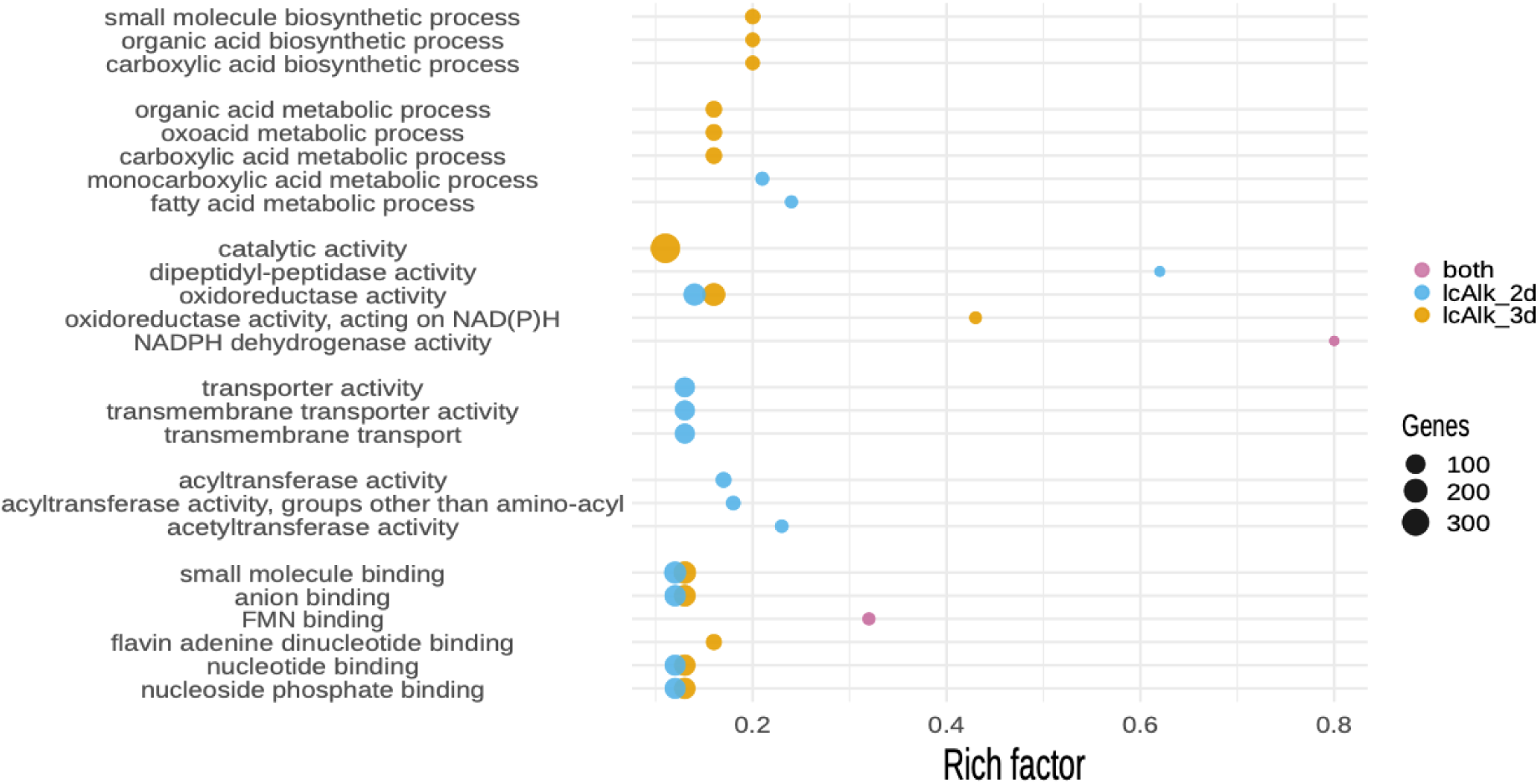
Bubble plot of gene ontology (GO) terms enriched in lcAlk_2d (orange) or lcAlk_3d (blue) vs C16_3d. Rich factor is the % of the transcribed genes of this GO term that are upregulated vs C16_3d. Bubble size is proportional to the number of upregulated genes.

Alkane metabolism involves their conversion to alcohols and carboxylic acids, and their subsequent catabolism through the well-studied fatty acid metabolism / β-oxidation pathway (Montazer et al., 2020). Two relevant, enriched processes are monocarboxylic acid metabolism and its child term, fatty acids metabolism (Fig. 3). Their upregulated genes include polyketide and fatty acid synthases, fatty acid desaturases, dehydrogenases, kinases and other transferases. Another interesting, enriched function is anion binding, particularly the binding of flavin mononucleotide (FMN), since many oxidases involved in alkane and fatty acid oxidation are flavoenzymes. Among these are the previously mentioned bacterial lcAlk hydroxylases LadA and AlmA (Feng et al., 2007; W. Wang & Shao, 2014). The genome of *A. parasiticus* encodes five luciferase-like monooxygenases with more than 40 % amino acid identity with LadA, consistent with finding in *A. flavus* (Perera et al., 2022) and one of them (RU639_010847) was upregulated in lcAlk_2d compared to C16_3d. Similarly, one of the seven genes classified within the same flavin monooxygenase family (IPR051820) as AlmA was upregulated in both lcAlk days.

GO enrichment in lcAlk_3d compared to C16_3d mirrored the enrichment of binding functions and oxidoreductase activities as in lcAlk_2d but exhibited unique differences. Notably, transmembrane and transport activities seen in lcAlk_2d were absent, replaced by an increase in metabolic and biosynthetic processes related to carboxylic acids and other small molecules. Of the 26 upregulated genes involved in carboxylic acid biosynthesis, 14 were also upregulated in lcAlk_2d. The accumulation of stable carboxylic acids results from the further oxidation of the generated alcohols, aldehydes and ketones during alkane metabolism (Hakkarainen & Albertsson, 2004). The use of alkanes for both energy generation and the biosynthesis of molecules such as lipids has been documented in entomopathogenic fungi (Napolitano & Juárez, 1997). In conclusion, the induction of transmembrane and transport activities needed for alkane internalization during the early response (lcAlk_2d) is followed by their incorporation into the metabolic processes of *A. parasiticus* MM36, as reflected in the shift towards biosynthetic and metabolic activities in lcAlk_3d.

The analysis of genes upregulated when the fungus was grown on C16 displayed a different profile. Although 1,800 genes were upregulated compared to lcAlk_2d, their high heterogeneity led to no enriched GO terms. However, when compared to lcAlk_3d, C16_3d exhibited strong enrichment of hydrolase activities, highlighting an abundance of glucoside hydrolases involved in carbohydrate metabolism, as well as peptidases, amidases, and esterases. While the strong enrichment of peptidases and amidases lacks a clear explanation, the esterases can be linked to the action of Baeyer-Villiger monooxygenases (BVMOs). BMVOs catalyze the conversion of a ketone to an ester, a reaction useful in case of sub-terminal alkane oxidation. Two BMVOs were upregulated in C16_3d compared to both lcAlk timepoints, while the other five displayed consistently low expression across all conditions. All seven BMVOs of *A. parasiticus* MM36 have orthologs in *A. flavus* with more than 94 % sequence identity, but these orthologs have demonstrated activity only up to n-C12 (Ferroni et al., 2014). This suggests that sub-terminal oxidation is more likely to occur in C16_3d rather than lcAlk, differing from the bacterium *Thalassolituus oleivorans* MIL-1, which uses terminal oxidation for C14, but shifts to sub-terminal oxidation for longer alkanes (Gregson et al., 2018).

### 2.6 Proposed mechanism of alkane assimilation by A. parasiticus MM36

Integrating the data from transcriptomic and secretomic analyses, we propose a potential mechanism for lcAlk functionalization, uptake and assimilation by *A. parasiticus* MM36 (Fig. 4). A secreted hydrophobin, a cerato platanin and three HsbA may act as surfactants to facilitate the initial enzyme and cell sorption to lcAlk. A variety of secreted enzymes that could oxyfunctionalize the alkanes is being secreted. Lignin active enzymes, including 4 MCOs and two peroxidases, can act in an unspecific manner potentially generating various oxygenated products and causing chain scission. Oxygenases including an UPO, four flavin-dependent oxygenases and a CYP505 hydroxylase can introduce terminal, sub-terminal, or di-hydroxyl groups into the alkanes, initiating their transformation into more reactive forms. These reactions are likely facilitated by ROS, which provide the necessary oxygen atoms. To mitigate potential ROS-related damage, *A. parasiticus* MM36 appears to produce ROS-controlling enzymes, including superoxide dismutases and catalases. Alcohols generated through these reactions could be further converted to aldehydes or ketones by five alcohol dehydrogenases, while three aldehyde dehydrogenases can convert the aldehydes to carboxylic acids. The secreted surfactants may interact with the produced hydrocarbons and facilitate cell adhesion and subsequent uptake through the cell wall and the membranes. This uptake could involve ABC and other membrane transporter families.

**Figure 4:**
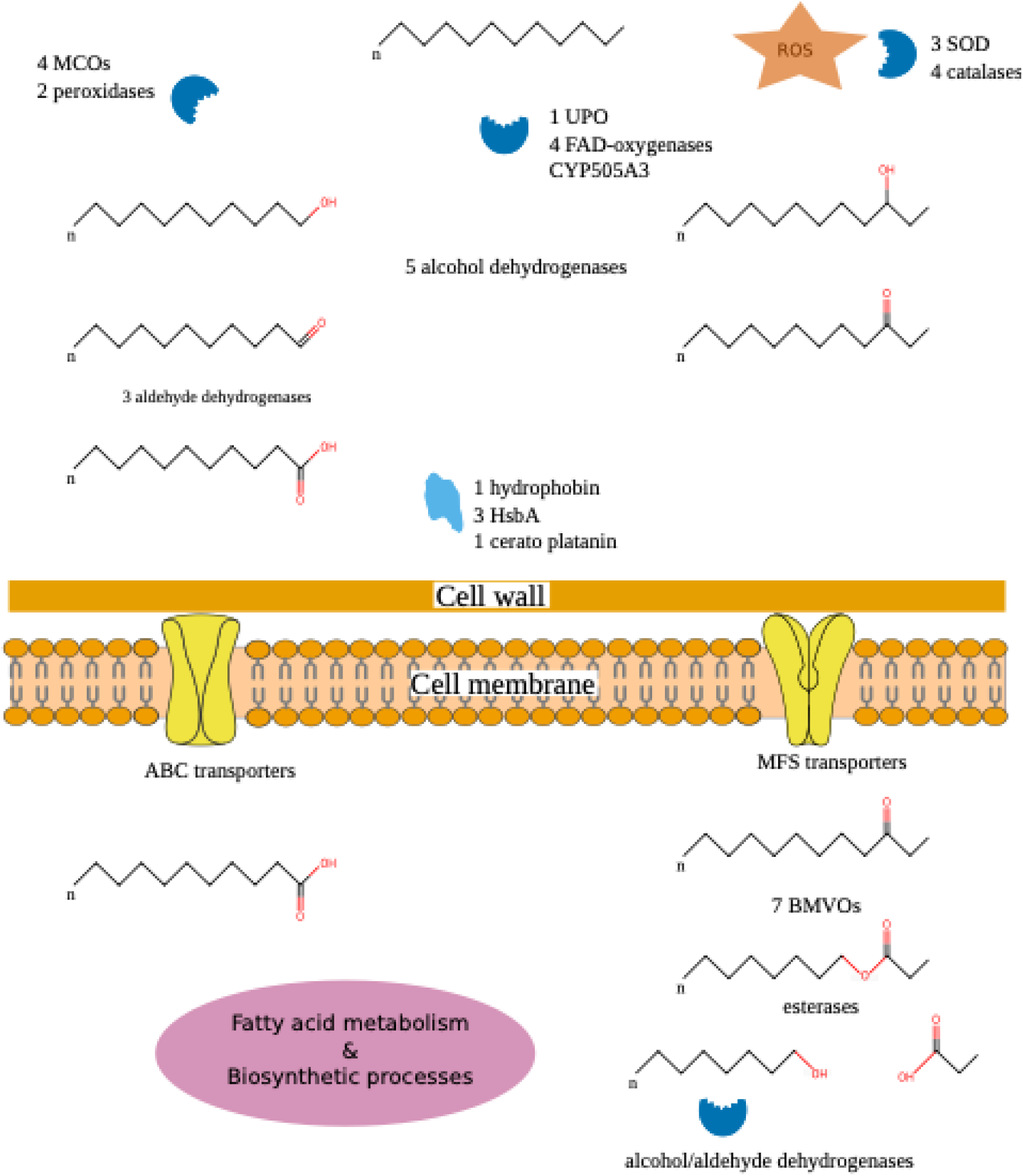
Proteomics- and transcriptomics-informed potential mechanism of alkane assimilation by *A. parasiticus* MM36. ROS: reactive oxygen species; MCO: multicopper oxidase; UPO: unspecific peroxygenase; SOD: superoxide dismutase; HsbA: hydrophobic surface-binding protein A; BMVO: Baeyer-Villiger monooxygenase.

Once inside the cell, these compounds can be directed toward catabolic or biosynthetic processes. In case of sub-terminal oxidation, the seven intracellular BMVOs may catalyze the conversion of ketones into esters, which could be hydrolyzed by a cascade of intracellular esterases, resulting in smaller chain carboxylic acids. The carboxylic acids are likely directed to the well-studied β oxidation pathway for energy generation or to biosynthetic pathways for lipid and small molecule production.

## 3. Materials and Methods

### 3.1 Genome sequencing and analysis

A. *parasiticus* MM36 was grown on YPD (1 % w/v yeast extract, 2 % w/v dextrose, 2 % w/v peptone) media plates supplemented with ampicillin sodium salt (Sigma, MI, USA) at a final concentration of 100 μg/mL. After sufficient growth was observed, spores were scraped from the agar plates and used to inoculate 100 mL YPD-ampicillin liquid cultures. Genomic DNA was extracted from 60 mg fungal biomass, obtained from a 2-day liquid culture at 27 °C, with shaking at 100 rpm. The biomass was filtered under vacuum in Miracloth, and cells were lysed using bead beading in a Tissue Lyser II instrument for 10 minutes at 25 Hz. The instructions of Quick-DNA Fungal/Bacterial Microprep Kit (ZymoResearch, CA, USA) were followed, using Lysis buffer with β-mercaptoethanol added, as suggested, to extract the DNA, dissolved in 10 mM Tris, pH 8.5, 0.1 mM EDTA.

A 350-bp insert size library was prepared and sequenced in paired-end mode (read length, 150 bp) by Novogene Europe on a NovaSeq 6000 instrument. Adapter sequences were removed (at an alignment score above 7, allowing 2 mismatches), and bases with a quality score below 10 and below an average of 20 on a 5-bp window were trimmed using Trimmomatic v.0.39 (Bolger et al., 2014). Reads smaller than 50 bases or with no pair (singletons) were discarded. Reads mapping to viral and bacterial genomes were removed with Kraken2 v.0.8 (Wood et al., 2019), with a confidence score of 0.7. *De novo* genome assembly was performed with Spades v3.15.4 with the isolate parameter (Prjibelski et al., 2020). Genome completeness was assessed with BUSCO v5.1.2 using the *Eurotiales* phylum single-copy orthologs (Manni et al., 2021).

Repeats were de novo identified with RepeatModeler v2.0 and LTRharvest (Ellinghaus et al., 2008) and were then searched against SwissProt after excluding transposases with an E-value threshold of 1e-10. Repeats aligned with protein-coding genes are removed with ProtExcluder v1.2. The genome was soft-masked with RepeatMasker v4.1.2 using the library of de novo identified repeats and the fungal repeat elements of Repbase 2017 and 2019 (Tarailo-Graovac & Chen, 2009).

Gene prediction was performed with BRAKER v3.0.3 (Gabriel et al., 2024) using the fungal partition of orthoDB v11 (Kuznetsov et al., 2023) and mapped RNAseq reads. The predicted genes and isoforms were reduced with TSEBRA (Gabriel et al., 2021) and the gene models were supplemented to MAKER v2.31.11 (Holt & Yandell, 2011) along with a Trinity assembly (Haas et al., 2014) of the RNA-Seq data to append the untranslated regions (UTRs) to the BRAKER gene models.

Proteins were functionally annotated with UniFIRE v2023.3. Gene and product names were assigned via UniFIRE and BLAST to SwissProt after filtering the alignments with Haas et al criteria (Haas et al., 2011). CAZymes were identified with run_dbcan v4.1.4 (Zheng et al., 2023). HMMER3 alignments were filtered with an e-value of 1e^−17^ and coverage 45 %. Proteins annotated only with DIAMOND were rerun in ultra-sensitive mode and filtered for 95 % coverage of both the query and subject sequence following Drula et al (Drula et al., 2022). Transporters were searched with BlastP and hmmsearch against TCDB downloaded in January 2024 (Saier Jr et al., 2021). Blast alignments were filtered for 70 % coverage and 30 % percent identity and hmmsearch for bitscore 100. Extracellular proteins were predicted with UniFIRE and DeepLoc2 (Thumuluri et al., 2022) or SignalP6 (Teufel et al., 2022) in the lack of transmembrane helices in the mature protein. Proteins with a signal peptide were searched for glycosylphosphatidylinositol (GPI) anchoring signal with NetGPI v1.1 (Gíslason et al., 2021).

### 3.2 Phylogenetic analysis

To classify *A. parasiticus* MM36 at the species level, all available proteomes included in a previous phylogenetic analysis of the *Aspergillus Flavi* section (Kjærbølling et al., 2020) were retrieved from Mycocosm (Grigoriev et al., 2014). Additionally, the reference genome of *A. sojae* SMF134 was downloaded from GenBank (accession GCA_008274985.1) and genes were predicted with GeneMark-ES v4.71 adapted for fungal genomes (Ter-Hovhannisyan et al., 2008).

Steenwyk et al (Steenwyk et al., 2024) generated profile Hidden Markov Models (HMMs) for single-copy orthologs of the *Aspergillus* genus. These HMMs were filtered for a length between 100 and 1000 amino acids and then 200 HMMs were randomly subsampled. The resulting HMMs were used to identify single-copy orthologs in the proteomes using orthofisher v.1.0.5 (Steenwyk & Rokas, 2021) and orthologs were aligned with mafft v7.490 (Katoh & Standley, 2013). Maximum-Likelihood phylogenetic analysis was performed with IQ-TREE v2.3.6 with an edge-linked fully-partitioned model (Minh, Schmidt, et al., 2020). Node support was calculated using 1,000 ultrafast bootstraps and 1,000 iterations of the SH-like approximate likelihood ratio test (Hoang et al., 2018). We calculated gene (gCF) and site concordance factors (sCF) by comparing single-gene trees to the concatenated tree (Minh, Hahn, et al., 2020).

### 3.3 Induction of PE-transforming enzymes and sample preparation for multi-omics analyses

Induction of potential PE-transforming enzymes was carried out in liquid cultures using mineral medium (MM) either supplemented with C16 or a lcAlk mixture. The growth media were prepared following the protocol described in a previous study (Taxeidis et al., 2023). Cultures were incubated at 27 °C, under continuous stirring at 120 rpm for 4 days. Each day, the mycelial biomass and the culture supernatant were collected separately. The fungal biomass was harvested under vacuum and washed with ultrapure water under aseptic conditions and immediately flash-frozen using liquid nitrogen. The culture supernatant was dialyzed overnight against 20 mM Tris-HCl buffer (pH 7.0) and freeze-dried under vacuum. After selecting the optimal time points the respective biomass samples were sent to Novogene B.V. (Netherlands) for RNA extraction, while corresponding supernatants were sent to the VIB Proteomics Core facility (Ghent, Belgium) for peptide purification and LC-MS/MS analysis. Both analyses were performed using triplicate biological samples.

The optimal time points for transcriptomics and proteomics analysis were determined through supplementary experiments aimed at identifying the time at which secreted enzymes effectively modify LDPE structure. Specifically, following the methodology detailed in a previous study (Taxeidis et al., 2023), each of the culture supernatants collected daily was tested for its ability to degrade LDPE powder. In this procedure, 50 mg of LDPE powder was incubated with 50 mL of the culture supernatant at 30 °C with continuous stirring at 160 rpm for 4 days. For control samples, the supernatant from MM/C16 cultures was used. After incubation, the plastic powder was removed, washed with 2 % (w/v) sodium dodecyl sulfate (SDS), and dried. The properties of LDPE were then analysed using Attenuated total reflectance-Fourier transform infrared spectroscopy (ATR-FTIR) to detect changes in the polymer’s molecular structure, following the same protocol as those mentioned in a previous study (Taxeidis et al., 2023).

### 3.4 LC-MS/MS analysis of the secretomes

Freeze-dried samples were dissolved in 5 % SDS in 100 mM triethyl ammonium bicarbonate (TEAB) to a final concentration of 1 µg protein / µL. From each sample 100 µg protein was reduced and alkylated by addition of 10 mM TCEP and 40 mM chloroacetamide (CAA). Phosphoric acid was added to a final concentration of 1.2 % and subsequently samples were diluted 7-fold with binding buffer containing 90 % methanol in 100 mM TEAB, pH 7.55. After centrifugation for 30 s at 4,000 x *g*, the columns were washed three times with 400 µL binding buffer and trypsin (1/100, w/w) was added for digestion overnight at 37 °C. Peptides were eluted in three times and dried completely by vacuum centrifugation. Peptides were re-dissolved in 100 µL loading solvent A (0.1 % TFA in water/ACN (98:2, v/v)) and pH was adjusted with 1 µL of 100 % TFA to a pH of 3. Samples were desalted on a reversed phase (RP) C18 OMIX tip (Agilent). After peptide binding peptides were eluted twice with 100 µL elution buffer (0.1 % TFA in water/ACN (40:60, v/v)). The combined elutions from each sample were dried in a vacuum concentrator.

Peptides were re-dissolved in 20 µL loading solvent A (0.1 % TFA in water/ACN (98:2, v/v)) of which 1 μL was injected for LC-MS/MS analysis on an Ultimate 3000 RSLCnano system in-line connected to a Q Exactive HF Biopharma mass spectrometer (Thermo). The peptides were separated on a 250 mm Aurora Ultimate, 1.7µm C18, 75 µm inner diameter (Ionopticks) kept at a constant temperature of 45 °C. The mass spectrometer was operated in data-dependent mode, automatically switching between MS and MS/MS acquisition for the 12 most abundant ion peaks per MS spectrum. Full-scan MS spectra (375-1500 m/z) were acquired at a resolution of 60,000 in the Orbitrap analyser after accumulation to a target value of 3,000,000. The 12 most intense ions above a threshold value of 15,000 were isolated with a width of 1.5 m/z for fragmentation at a normalized collision energy of 28 % after filling the trap at a target value of 100,000 for maximum 120 ms. QCloud has been used to control instrument longitudinal performance during the project (Chiva et al., 2018).

### 3.5 Processing of the LC-MS/MS data

The raw data were searched together using the nf-core/quantms v1.3.0dev (Dai et al., 2024) pipeline of the nf-core collection of workflows (Ewels et al., 2020) against the proteome of *A. parasiticus* MM36 and a proteomics contaminant library (Frankenfield et al., 2022). The pipeline was executed with Nextflow v24.04.2 (Di Tommaso et al., 2017), utilizing reproducible software environments from the Bioconda (Grüning et al., 2018) and Biocontainers (da Veiga Leprevost et al., 2017) projects. Precursor mass tolerance was set to 15 ppm based on Param-Medic (May et al., 2017) and fragment mass tolerance was set to 0.02 Da. Trypsin cleavage was allowed a maximum of two missed cleavages. The peptide identification step was performed with Comet, MS-GF+ and SAGE (Eng et al., 2013; Kim & Pevzner, 2014; Lazear, 2023), with search engine results rescored by MS2Rescore (Declercq et al., 2022). Variable modifications were set to oxidation of methionine and tryptophan, acetylation of protein N-termini, deamidation of asparagine and N-terminal glutamine conversion to pyroglutamic acid and ammonia loss of cysteine. Carbamidomethylation of cysteine residues was set as a fixed modification. These modifications were determined after an open search of the raw data using Fragpipe with default settings (Geiszler et al., 2021; Kong et al., 2017).

The peptide level intensities were imported in the prolfqua package (Wolski et al., 2023), log2 transformed and robust z-score scaled. Protein intensities were estimated from peptide intensities using Tukey’s median polish (TMP). However, TMP summarization creates artifacts in cases where all peptides of a protein are detected uniquely in single samples, yielding misleading uniform protein intensities across all samples. This phenomenon is previously reported in microarray data summarization (Giorgi et al., 2010). For proteins affected by this artifact, median summarization of the top three peptides was employed instead. Subsequently, only reliably quantified proteins present in at least two biological replicates of one condition were retained for downstream analysis. Differential protein abundance was tested using the empirical Bayes approach with imputation of missing values as implemented in prolfqua.

### 3.6 Transcriptome analysis during alkane assimilation

Total RNA was extracted from each timepoint in triplicates by Novogene, the polyA+ fraction was purified, and libraries were constructed with the Novogene NGS RNA Library Prep Set (PT042)* kit and sequenced in paired-end mode on a NovaSeq 6000. One C16 sample with a RIN value of 3.7 was discarded leaving this condition with two replicates. Raw data were processed using nf-core/rnaseq v3.10.1 (Patel et al., 2024) of the nf-core collection of workflows (Ewels et al., 2020) at the high-performance computing bioinformatics platform of HCMR (Crete, Greece) (Zafeiropoulos et al., 2021). Raw counts were loaded in edgeR v3.43.4 and were normalized using the TMM normalization (Robinson et al., 2010). Based on Chen et. al. (Chen et al., 2016) genes with low expression were excluded. Specifically, only genes with more than 10/L reads, where L=16.66 represents the median library size in millions, in at least two replicates of any condition were retained for downstream analysis. Differential expression (DE) was tested with the edgeR-exact test following recommendations for few replicates (Schurch et al., 2016). The p-value was adjusted with the Benjamini-Hochberg (B-H) correction and corresponds to an FDR of 5 %. Enrichment of gene ontology (GO) terms was tested with goatools v1.3.1 (Klopfenstein et al., 2018) with the go-basic.obo release 2023-04-01 using Fisher’s exact test with 5 % B-H FDR. GO terms were derived from the Interproscan annotations.

#### Data availability

The Whole Genome Shotgun project has been deposited at DDBJ/ENA/GenBank under the accession JAWDVE000000000. The transcriptomic RNA-seq data have been deposited to the NCBI Gene Expression Omnibus (GEO) database with the dataset identifier GSE282836. The mass spectrometry proteomics data have been deposited to the ProteomeXchange Consortium via the PRIDE (Perez-Riverol et al., 2022) partner repository with the dataset identifier PXD058271 and 10.6019/PXD058271. Code for the transcriptomic and proteomic analysis is available in GitHub (https://github.com/Roman-Si/Asp-parasiticus_alkane_multiomics).

## Funding

This research was funded by the Hellenic Foundation for Research and Innovation (H.F.R.I.) under the “2nd Call for H⋅F.R.I. Research Projects to support Faculty Members and Researchers” (Project Number: 03061, PlastOmics).

## Acknowledgments

The authors would like to thank the VIB Proteomics Core for the contribution regarding the mass spectrometry-based proteomics experiments (EPIC-XS, project number 823839, funded by the Horizon 2020 programme of the European Union). This research was supported in part through computational resources provided by IMBBC (Institute of Marine Biology, Biotechnology and Aquaculture) of the HCMR (Hellenic Centre for Marine Research). The authors thank Dr Anastasia Gioti for extracting DNA from *A. parasiticus* MM36 and Professor Stamatina Vouyiouka for providing equipment for the Attenuated total reflectance-Fourier transform infrared spectroscopy (ATR-FTIR) analysis.

